# RCaN : a software for Chance and Necessity modelling

**DOI:** 10.1101/2021.06.09.447734

**Authors:** Hilaire Drouineau, Benjamin Planque, Christian Mullon

## Abstract

Uncertainty is a challenge in modelling ecological systems and has been a source of misunderstandings between modelers and non-modelers. The “Chance and Necessity” (CaN) modelling approach has been proposed to address this issue, in the case of trophic network modelling. CaN modelling focuses exploring food-web trajectories that can satisfy fundamental physical and biological laws, while being compatible with observations and domain knowledge. This type of approach can facilitate discussion among actors as it promotes sharing of information and does not presuppose any knowledge of modelling practices. It is therefore suitable for participatory modelling, i.e. a modelling approach in which different actors can confront their knowledge and understanding of the marine system and of the associated uncertainties.

One important ingredient to achieve participatory modelling is the availability of a modelling platform that is efficient, fast and transparent, so that all actors can understand and follow the modelling steps and choices, and can rapidly visualize and discuss the results. But, until now, there existed no software to easily perform CaN modelling. Here, we present RCaN and RCaNconstructor. Combined, these provide the first tool to build CaN models in an intuitive way that is 1) suitable within participatory frameworks, 2) transparent, 4) computationally efficient, 5) fully documented through the provision of meta-information and 6) supportive of exploratory analyses through predefined graphical functions.

## 1 Introduction

Modelling complex ecological systems is often a trade-off between the accurate representation of the many components and interactions within an ecosystem and the reduction of ecosystem processes to few and important elements. Rather than attempting to model multiple ecological processes in detail, CaN modelling focuses on exploring possible ecological dynamics given a set of constraints. The principles of CaN modelling and the corresponding mathematical formulation have been outlined in [1]. The general idea is that it is easier for various actors (managers, scientists, stakeholders) to agree about ecological constraints, which separate what is possible from what isn’t, than to agree on the mechanistic formulation of detailed ecological processes. This is particularly true when modelling complex ecological systems for which available information/observations are limited. This approach is called “Chance and Necessity” modelling (CaN): chance refers to the indeterminacy of ecological processes while necessity reflects the constraints that delineate what is possible from what isn’t.

While the CaN principles can be applied to a wide range of ecological problems, the first application was specifically designed to explore the temporal dynamics of marine systems [1]. This has been motivated by the shift from conventional single-species fisheries management towards ecosytem-based fishery management (EBFM, see for example [2, 3]). This shift implies up-scaling of the traditional population scale management into a complex set of biotic (e.g. trophic food interactions) and abiotic sets of interactions (e.g. interactions with physical habitats). This has led to the development of complex ecological models, but which ability to help management is questioned by many actors who may have limited trust in such models [4]. In this context, there is a call for transparency in the science that supports management, and participatory modelling and assessment have been proposed as a way to support mutual trust. CaN is proposed as one tool to support transparency and participatory modelling by providing a simplified food-web modelling framework, in which the ideas of uncertainty and ecological limits are central. Assumptions about knowns and unknowns and about possible and impossible processes are presented explicitly and all model constituents can be described in plain language (little or no modelling jargon).

By contributing to the transfer of biomass, energy and pollutants in ecosystems, trophic interactions play a major role in ecosystem functioning [5]. Trophic food web models have proved to be key tools for the quantification of these fluxes, they provide a holistic view of ecosystem functioning, and are increasingly used to assess ecosystem ecological status and to explore the effect of anthropogenic pressures such as fisheries [6]. Food web models, mostly Ecopath with Ecosim (EwE) [7, 8], have played an important role in the implementation of the EBFM. CaN modelling further contributes to this effort by providing a modelling framework for marine food-webs that primarily relies on ecological constraints which can be agreed upon by diverse actors.

While the equations behind CaN modelling are simple, one major technical challenge is to sample a polytope in a highly dimensional space. This is particularly useful if achieved inside the R framework [9], which is widely used by ecological modelers.

Until now, there existed no software to easily perform CaN modelling. The first models were constructed in an *ad hoc* fashion by assembling and structuring relevant data and developing tailor-made programs in Mathematica or R to run the model. This made it difficult to communicate and replicate CaN models. It also restrained the access of the model to non-modelers. Making CaN modelling transparent, replicable and efficient in a participatory modelling framework requires that model construction, sampling and output visualization be easily accessible by modelers and non-modelers alike, that the structure of CaN models be standardised and that the model sampling be robust and efficient. A graphical user interface (GUI), which provides an easy entry level for users is also a highly desirable feature for a modelling approach targeting non-modelers. In this contribution, we present a software for the implementation of CaN modelling in R, called RCaN, and an associated GUI in Java, called RCaNconstructor. RCaN and RCaNconstructor can be used jointly or separately. Their combined use allows users to easily construct, document, sample and interpret the results of CaN food-web models. The relationship between RCaN and RCaNconstructor is shown in figure 1. The main features of RCaN and RCaNconstructor include:

1. a user-friendly interface that allow multiple users to jointly develop and revise models. Users can then focus on the discussion about their knowledge of the ecological system and how to best structure a model that fits their needs,
2. a file and data management system that ensures consistency between the different elements of a CaN model. For example, it must be guaranteed that each link connects existing components or that each constraint points to components, links or data series that are adequately specified in the model,
3. a standardized file format to fully specify a CaN model,
4. a Graphical User Interface that facilitate the handling and replicability of data use and the tracking of observations sources (data files),
5. a set of R functions that allow experimented programmers to construct, manipulate and visualize the outputs of CaN models in a flexible manner,
6. standardized graphical outputs of the model results, which promote discussions among modelers and other actors and ease the interpretation of model results,
7. a system for registering meta-information about the model, such as model version, authors, data sources, assumptions, and so on. This ensures transparency and reproducibility of modelling experiments.

**Figure 1:**
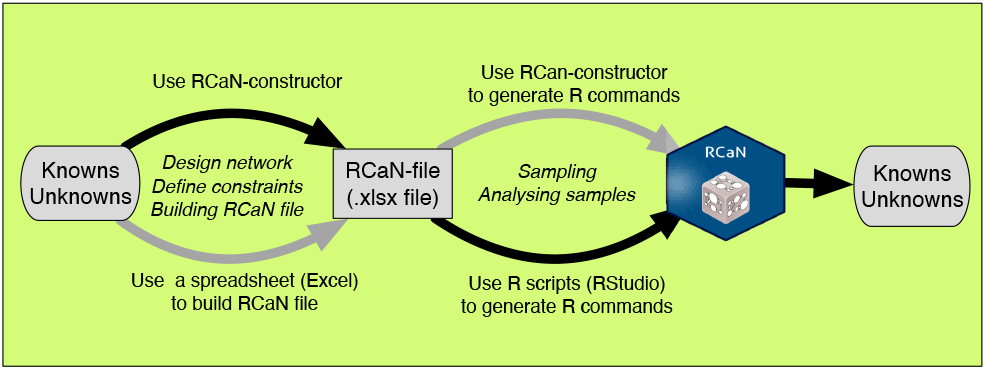
The articulation between RCaN and RCaNconstructor with two main stages, (a) building files and (b) building a polytope, sampling it and analyzing results. There are several ways to go through these stages; the ones that have been followed for the Barents Sea example (section 6) appear in dark gray in the figure. RCaN and RCanConstructor have been designed for being part of a participatory approach: the design of the file, especially of constraints, and the interpretation of most results are simple and easy to understand.

In section 2 we summarize the elements of the CaN modelling approach. In section 3, we present the format of an RCaN file. In section 4 we present the R library RCaN which is designed to construct, sample and produce graphical outputs of CaN food-web models. In section 5 we introduce RCaNconstructor, the graphical user interface that facilitates the co-construction of CaN models and the inclusion of meta-information. In section 6, we briefly describe the Barents Sea trophic network which we use as a case study example. Finally, we discuss the possible future developments of the RCaN and RCaNconstructor softwares.

## 2 Principles of the CaN approach

In CaN, food-web dynamics is defined by time-trajectories of biomass and fluxes between trophic groups (also termed components). The biomass of all trophic groups and the fluxes between them can usually not be precisely observed. Therefore, the food-web dynamics is indeterminate, and a range of dynamics is possible. However, not everything is possible because some ecological or physiological constraints are known to bound the temporal evolution of ecological system, and/or because some observations can inform about the past dynamics of some components of the food-web. These eco-physiological and observational constraints can be translated into mathematical constraints that need to be satisfied before a food-web trajectory can be considered possible. The ensemble of these mathematical constraints constitutes the “necessity” in a CaN model. It is by focusing on constraints that can be easily expressed and communicated between actors and by explicitly acknowledging indeterminacy in natural systems that CaN modelling facilitate exchanges among actors to support resource management.

### 2.1 Network structure

In CaN modelling, the food-web is defined as a network. In this network, each vertex/node is called a component which can be a species, a trophospecies, i.e., a group or sub-group of species that share common prey and predators, or another component exchanging biomass with the rest of the systemt (e.g. a fishery). Components may be included within the model domain, in which case their biomass trajectories are explicitly considered. Alternatively, they can be outside the model domain, in which case it is only the fluxes from/to these components that are considered, but not the components trajectories. An edge/link corresponds to a flux of biomass between two components. Most fluxes are trophic: from a prey to a predator. Some fluxes are non-trophic, such as import/export of biomass from adjacent regions or removal of biomass through fishing.

### 2.2 Components’ characteristics

Components inside the system are characterized by the following biological properties. **Satiation** reflects the maximum consumption rate per unit biomass of the predator. **Inertia** expresses that variations in biomass from one year to the next are bounded between a maximum growth rate and a maximum mortality rate. **Digestibility** is a correction factor that accounts for variations in energy content between prey. **Assimilation efficiency** expresses the proportion of the biomass ingested by a predator that is effectively assimilated; the product of the potential assimilation efficiency by the digestibility correction factor is the absorption efficiency (the proportion of prey biomass digested and absorbed). **Refuge biomass** is the absolute minimum biomass that a trophospecies can reach. **Other losses** is a mortality coefficient that account for losses, i.e., metabolic losses and other mortality, not explicitly accounted for in the model.

Components outside the model domain do not require input parameter except when they constitute a prey, in which case digestibility should be provided.

### 2.3 Observations

CaN modelling builds from past observations of the dynamics of the food-web. Observations can be catch or landings data, biological samples of specific populations, stomach samples or other sources of diet information, survey abundance estimates, outputs from stock assessment models, estimated ranges of biomass, and so on.

### 2.4 Constraints

Constraints that limit the dynamics of the food-web are at the heart of CaN modelling and they are the way by which most of the knowledge and data enter a CaN model. Some constraints are implicitly incorporated into CaN while others can be explicitly formulated.

CaN models assume conservation of energy during trophic exchanges, as do many other trophic models. A set of implicit constraints is standard for all CaN models: (a) biomass of compartments are positive and above refuge biomass, (b) trophic fluxes between components are positive, (c) the sum of fluxes into a compartment is limited according to its biomass (satiation), (d) the proportional changes in biomass of a compartment are bounded (inertia).

Additional explicit constraints can be provided. These are written in the form of inequalities or equalities. The left side of the (in)equality must contains a reference to one or several compartments or fluxes. The right side of the (in)equality can contain fixed values, compartments, fluxes and observational data-series.

There are two kinds of additional explicit constraints. Some constraints are the expression of a knowledge about system functioning and relate the biomass of a component to its incoming or outgoing fluxes. Other constraints restrict the possible values of a biomass or a flux on the basis of available observations. These last constraints constitute the way to deal with observation uncertainties in CaN modelling.

### 2.5 Trajectories

A food-web trajectory is defined by the ensemble of biomass fluxes at each time-step and by the initial food-web configuration, i.e., the initial biomass in each component. Each trajectory can be expressed as a single point in a multi-dimensional space of dimension *P* = *nF* × *nt* + *nB*_0_, with *nF* the number of fluxes, nt the number of time steps and *nB*_0_ the number of initial biomasses. The number of dimensions can be high. For example, following the dynamics of a food-web with 10 components and 20 fluxes over 15 time steps will results in *P* = 20 × 15 + 10 = 310 dimensions.

All constraints are linear equalities or inequalities that restrict the space of possible food-web trajectories. As constraints are linear, the set of the trajectories that satisfy all constraints has the shape of a convex polytope in a high dimensional space.

### 2.6 Sampling trajectories

The main objective of CaN modelling is to sample food-web trajectories that are possible given (1) the food-web structure, (2) the species biological properties, (3) available observations and (4) constraints. The diversity of sampled trajectories expresses the indeterminacy in the system.

Sampling food-web trajectories is achieved by sampling this high-dimension polytope using dedicated sampling algorithms.

## 3 RCaN file

In RCaN, the input information describing the model structure is provided in a spreadsheet in the xlsx format. In the worksheet **Components**, the first column corresponds to the name of a component. The second column indicates whether the component is inside our outside the model domain. Other columns correspond to components characteristics (i.e. input parameters) that have been presented in section 2.2).

In the worksheet **Trophic fluxes**, each flux is characterized by an Id (first column) and links a source component (column From) to a sink component (column To) and by its type, trophic or non trophic (last column); in a trophic flux, digestibility and assimilation should be applied to the flux.

In the worksheet **Observations**, the first column gives the identifier of time steps (most of the time years). Other columns correspond to the observational data-series that can be used to constrain the system (see 2.3).

In the worksheet **Constraints**, each row describes a constraint. Each constraint has an Id (first column), a formula defining the linear constraints between fluxes, biomass and observations (second column), a time period of validity (third column) and a flag indicating whether the constraint should be active or not (fourth column). The contents of the above worksheets are interrelated and naming of components and fluxes across worksheets should be consistent.

The additional worksheet **INFO** contains meta-information about the model. It is not compulsory to provide this worksheet to be able to run RCaN, but it is highly recommended to do so. The worksheet contains information about the model contributors, the model version, the source of data and information, etc., all of which contribute to transparency and replicability.

## 4 RcaN

### 4.1 Introduction

RCaN is an R package to build, sample and visualize the results of CaN food-web models. It does this by 1) constructing the polytope from implicit/explicit constraints, 2) solving the numerical challenges of polytope sampling and 3) producing graphical outputs. The next sections detail the technical features of the implementation of these different steps, in the R language.

### 4.2 The different steps to use RCaN

#### 4.2.1 Build a polytope from RCaN-file

Once the RCaN-file has been built, a first step is to organize the information and to build the polytope describing the model. More precisely, the mass conservation of biomasses and constraints can be summarized in matrix notation, and all these matrices should be automatically generated whatever the constraints and parameters used by the modeler.

Constraints can be summarized in a matrix form as *A* · *x* <= *b* (inequality constraints) and *C* · *x* = *v* (equality constraints) with x a solution (e.g. a trajectory defined by components’ biomass at initial step and fluxes at each time step), A and C two matrices with as many columns as parameters in the model (fluxes per time step and initial biomass per component) and as many rows as constraints. Vectors *v* and *b* specify the bounds of the equalities and inequalities.

Using the constraints formula and the symbolic links among variables, RCaN automatically builds matrices *C* and *A* and vectors *b* and *v* for each active constraint. This is done by a simple call to the function **buildCaN** which reads the information from the RCaN-file and returns a single object that contains the trophic network and the corresponding polytope matrices. For example, in the Barents Sea case study (section 6), the function automatically constructs the matrix A which has 2593 rows and 775 columns, and the matrix C which has 54 rows (and by definition, the same number of columns).

#### 4.2.2 Checking polytope characteristics

At first try, the specification of model can lead to an empty polytope (i.e., a model with no solution that satisfies all the constraints) or an unbounded polytope (i.e., a model in which the specified constraints are not sufficient to bound a given parameter that can vary subsequently to infinity). Before running the model, it is wise to check the status of the polytope (the model won’t run with an empty polytope while results are generally not relevant with unbounded polytopes). The package provides different functions to check the properties of the polytope, of which the two most important are **checkPolytopeStatus**, which checks whether the polytope is closed and non-empty, and **getAllBoundsParam**, which estimates the bounds for all parameters. If the model is empty, the RCaN function **findingIncompatibleConstr** can be used to identify which constraints are incompatible with each other.

#### 4.2.3 Sampling polytope

The aim of this step is to perform a uniform sampling within a high dimensional polytope. The function **sampleCaN** carries out the uniform sampling and returns results in the standard **coda** format of a **mcmc.list** [10]. It is possible to sample several chains in parallel.

#### 4.2.4 Diagnostics about sampling

Once sampling is done, it is necessary to check whether the uniform sampling was successful. Since the result of **sampleCaN** is a mcmc.list object, we can use all the diagnostics tools provided in the package coda, such **traceplot**, **summary** or **gelman.diag** or autocorrelograms.

#### 4.2.5 Graphical analysis of sampling

RCaN provides several functions to explore the model results. The function **ggSeries** plots the time-trajectories of biomass and fluxes and **ggViolin** plots their empirical distributions. The function **ggDiet** provides a summary of the diet fractions of different prey for each predator. **ggPairsBiomass** displays the pairwise relationships between species biomasses. The function **ggTrophicRelation** displays the relation between prey abundance and ingestion. **ggGrowth** plots annual growth against population biomass, which is useful to discuss density-dependence. **ggSatiation** shows how much a predator ingests against the predator biomass and displays the maximum possible feeding (satiation). **ggSatiatInertia** shows the standardized growth against standardized ingestion and is useful to investigate possible bottom controls. **ggBottelneck** aims to detect potential bottleneck species by showing whether a species was close from satiation (i.e., has more resources that it can assimilate) whereas its predators were far from satiation (i.e., its predators could potentially have eaten more if food was available).

### 4.3 Technical considerations

The development of the packages raises several challenges. The first challenge was to improve computation performance. For this purpose, RCaN is interfaced with **C++** using two R packages **Rcpp** [11] and **RcppEigen** [12]. The second challenge was to check the characteristics of the polytope. This was achieved by using standard linear programming library: RCaN uses **lp_solve** through the R package **lpSolveAPI** [13]). The third challenge was also to translate the content of a RCaN-file into matrices, and especially to allow user to specify constraints in a simple language. This was made possible by using the R package **symengine** [14], an interface to the **symengine** symbolic manipulation library (the syntax of explicit constraints is presented in Appendix A.

RCaN can performs uniform sampling with two methods, the hit-and-run and the Gibbs sampler. Hit-and-run is a well-known algorithm for uniform sampling of high-dimensional convex space [15][16]. Basically, hit-and-run sampling starts from a point in the polytope. Then at each iteration, a direction of the high dimensional space is randomly drawn, and a new point is uniformly sampled between the two extreme bounds defined by the intersection with the envelop of the polytope. At each iteration, the point is moved in a single direction of the multidimensional space, and consequently, when the sampler is “trapped” in a narrow zone of the polytope, it may take many iterations to pick the appropriate direction to exit the narrow zone resulting in autocorrelation. A variation of the hit-and-run sampler is referred to as the Gibbs sampler or “Coordinate Hit-and-Run” [16][17]: instead of picking a single direction and updating all coordinates in this direction, at each iteration the Gibbs sampling successively and independently updates the value of each parameter (i.e., at each iteration, the sampler moves successively in all directions of the space) to generate a new parameter value that satisfies the constraints. By doing so, it guarantees an efficient convergence [17]. By default, RCaN uses the more efficient Gibbs’s sampler.

## 5 RCaNconstructor

The main goal of RCaNconstructor is to help users to construct and revise CaN models in a participatory context. Internally, it ensures the consistency between the different worksheets of the RCaN-file (changing the characteristics of an object somewhere triggers the corresponding changes elsewhere). In addition, RCaNconstructor facilitates the elicitation and tracking of meta-information. RCaNconstructor can also execute RCaN commands.

RCaNconstructor is a Java program using Javafx for building the graphical interface and allowing the code to be imported and run on external computers; it uses the Java library **RCaller** for interfacing R and Java.

### 5.1 Meta information

Users can document the meta-information relevant for their model, using the RCaNconstructor Meta information menu. The meta information template invites contributions relevant to model version and authors, domain coverage and units, definition of components and fluxes, sources for input parameters and observations, rational for constraints, model uncertainties and limitations.

### 5.2 Views

The View menu opens a collection of graphical interfaces. With these, the user can graphically draw the food-web structure (View/Network) enter the input parameters (View/Components), edit trophic links (View/Fluxes), load observation files (View/Observations) and specify the model constraints (View/Constraints). The meta-information and the model setup are saved in a ready to use RCaN-file.

### 5.3 R interface

The RCaNconstructor is interfaced to R with the RCaller library [18]. From the RCaNconstructor menu, the user can interactively: (a) start a R session and load RCaN library, (b) Build the model polytope (corresponds to the **buildCaN** command of RCaN), (c) sample the polytope (corresponds to the command **sampleCaN**), (d) retrieve sampling diagnostics and (e) produce graphical outputs of the model results.

## 6 Example: the Barents Sea food-web

To illustrate CaN modeling using RCaN and RCaNconstructor, we use a simplified food-web of the Barents Sea as an example.

In what we present below, we have chosen to build the RCaN-file using RCaNconstructor and then to run RCaN commands to construct the polytope, sample it, and analyze the properties of the sample.

A more detailed presentation of the Barents Sea example is provided in the Supplementary material.

### 6.1 Monitoring of the Barents Sea

Monitoring of the Barents Ecosystem has been conducted for several decades and a collection of data series are reported regularly and compiled by the ICES working group on the integrated assessment of the Barents Sea (WGIBAR, [19]). These time series cover the period 1988-2019 and include data on:

1. landings for pelagic and demersal fish, krill, shrimps and marine mammals,
2. survey-based biomass estimates of zooplankton, pelagic and demersal fish,
3. satellite-based estimate of Net Primary Production (NPP, from 1998),
4. consumption estimates by Atlantic cod (*Gadus morhua*) of krill, shrimps, capelin (*Mallotus villosus*), herring (*Clupea harengus*), polar cod (*Boreogadus saida*) and Atlantic cod.

In addition, there are estimates of the minimum and maximum plausible biomasses of benthos, marine mammals and birds which are used as limits for the whole time-period in the model ([20]).

### 6.2 Meta Information

Information about the model version, authors, funding sources, data sources and pre-processing, assumptions and uncertainties are provided in the model RCaN-file.

### 6.3 Food-web Structure

The model of the Barents Sea food web is composed of 11 components. Four of them are external (two plankton standing stocks from the adjacent Norwegian Sea, primary producers and Fisheries), and seven are within the model domain, and include plankton (Hzoo: herbivorous zooplankton, Ozoo: omnivorous zooplankton), benthos (bent), fishes (PelF: pelagic fishes, DemF: demersal fishes) and top predators (MM: marine mammals, birds). These 11 components are connected via 24 biomass fluxes among which 18 correspond to trophic relationships and 6 to imports (plankton) or export (fisheries catches) of biomass.

With RCaNconstructor the food-web structure is specified by drawing the components and fluxes (menu View/Network). Using this graphical interface (fig. 2, top-left) it is possible interactively add, delete and edit components and fluxes. The graphical positioning of the components of the network can also be modified to ease the visual interpretation of the food-web structure.

**Figure 2:**
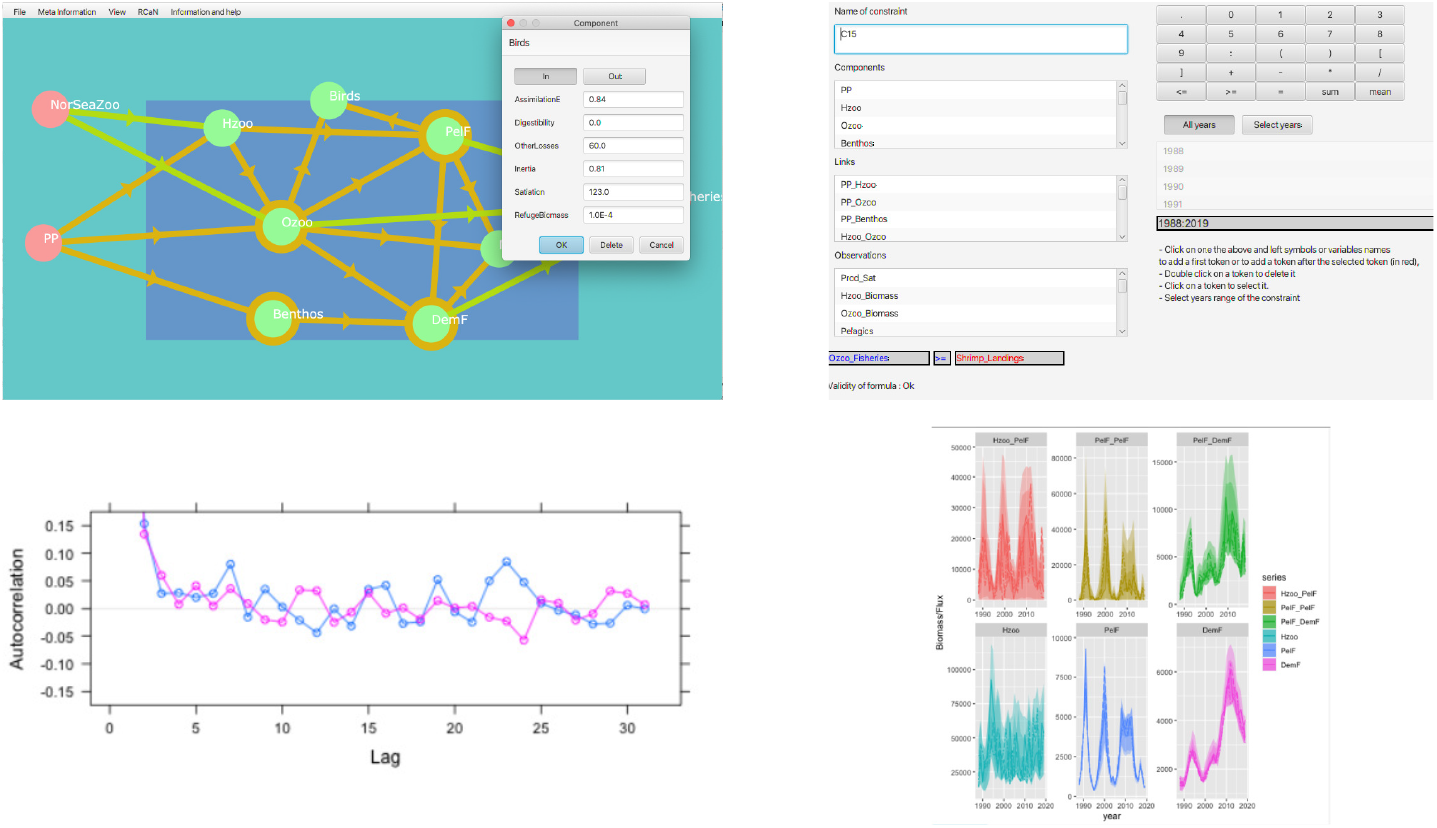
A selection of input/output windows from the RCaN-constructor applied to the Barents Sea food-web model. Top left: RCaNconstructor graphical interface to draw and specify the trophic network. Top right: the graphical interface used to specify the model constraints. Bottom left: sample diagnostic plot showing the autocorrelation of two chains of one flux. Bottom right: sampled trajectories (time-series) of selected fluxes and components.

### 6.4 Input parameters

The input parameters for individual components of the Barents Sea food web (menu View/Components) are taken from [20]. These were derived from the Metabolic Theory of Ecology, Life-history theory or previous modeling exercises using for example Ecopath [21]. These are summarized in Table 1, in the supplementary material.

### 6.5 Observations

The observation data series are derived from to the Barents Sea monitoring program (section 6.1). These includes time-series of biomass for all species groups within the model domain as well as information about some of the fluxes (e.g. fluxes from primary producers, fluxes towards fisheries, and fluxes between species).

### 6.6 Explicit constraints

In addition to implicit constraints (satiation, inertia, positive fluxes and refuge biomass), the Barents Sea RCan model includes explicit constraints. These constraints reflect additional knowledge about the ecological system and can be used to link the model to available observations. These are as follows:

1. The consumption of primary production by zooplankton and benthos is constrained by net primary production estimates (± 30%) for the period when these estimates are available (1998 to 2019). For other years, this sum is bounded by estimates of absolute maximum and minimum primary production that are set to 1/2 million and 2 million tons respectively.
2. Time-series of landings provide accurate information about the fluxes from herbivorous and omnivorous zooplankton, pelagic and demersal fish, and marine mammals to corresponding fisheries. Therefore, these fluxes can be set to the values of the estimated landings.
3. The annual estimates of biomass for herbivorous and omnivorous zooplankton, and pelagic and demersal fish, derived from research surveys, can be used to constrain the biomass of these groups.
4. The estimates of minimum and maximum plausible biomasses for benthos, marine mammals and birds can be used as limits for these groups for whole time-period in the model.
5. The food consumption by demersal fish can be bounded based on consumption estimates by cod in the Barents Sea. These bounds are as follows:
  a. the consumption of demersal fish by demersal fish is greater than the consumption of cod by cod,
  b. the consumption of pelagic fish by demersal fish is greater than the consumption of capelin, herring and polar cod by cod;
  c. the consumption of omnivorous zooplankton by demersal fish is greater than the consumption of krill and shrimp by cod
  d. the total consumption by demersal fish is not greater than twice the the total consumption estimates by cod.

### 6.7 Polytope building

Once the components, input parameters, fluxes, observations and constraints are provided, the model is fully specified. This information is gathered in the corresponding RCaN file. The polytope can be constructed by running R commands of the RCaN library. This can be achieved within the RCaNconstructor environment or directly within the R environment. Both ways have their advantages and limitations. Using RcaNconstructor is intuitive and does not require prior knowledge of the R language. Using R provides a greater flexibility for users who are familiar with the R language. We provide below the commands in the R language.

We build the polytope with following R command :

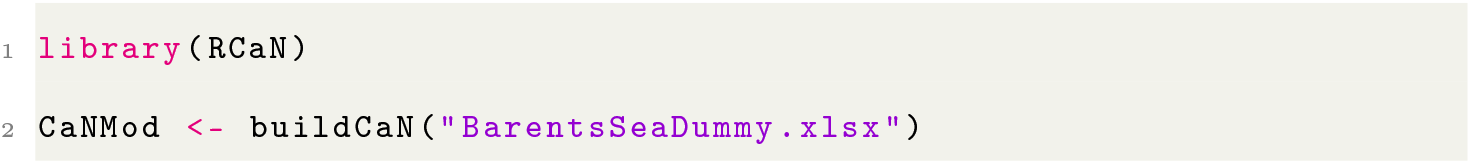

In the Barents Sea case, the polytope has 775 dimensions (7 initial values - one for each component within the model domain - and 24 fluxes times 32 years). There are 2593 constraints (both implicit and explicit). Constructing this polytope on a laptop computer takes about 70s (MacBook Pro Retina, 13-inch, Early 2015, 3.1 GHz Dual-Core Intel Core i7).

It is possible to check the status of the polytope (bounded, empty or unbounded) and to compute the bounds of the polytope in every of its 775 dimensions using the following commands (the time taken to obtain the results is indicated in round brackets):

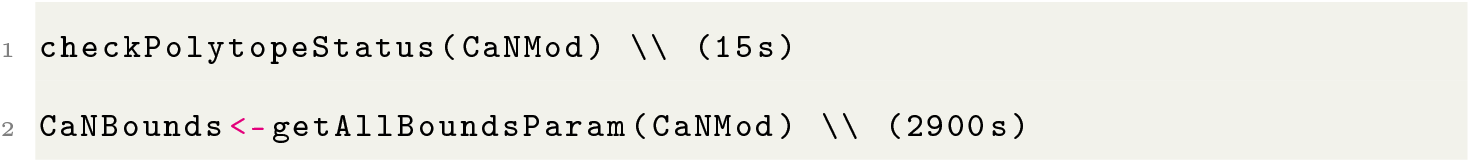

### 6.8 Polytope sampling

Once the polytope is built, and if it is properly shaped (i.e. not empty or unbounded), it is then possible to sample it. Each sample corresponds to a trajectory of the food-web. It contains the initial biomass values for each component inside the model domain and the values of the fluxes at each time step. That is 775 values in the example of the Barents Sea model. Sampling the polytope is achieved with one RCaN command:

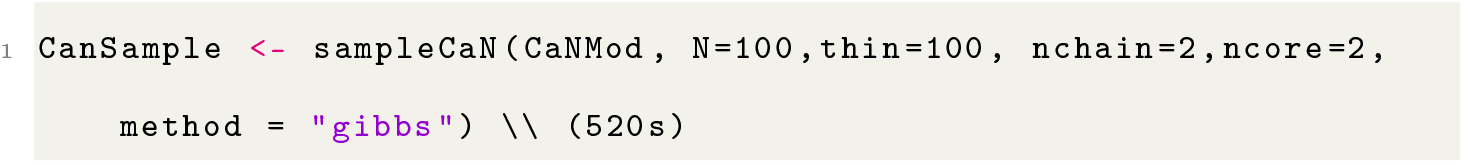

This command returns an R list object which contains the original model structure and two chains of 100 samples (i.e. 100 food-web trajectories). In total 10,000 samples were explored in each chain, but only every 100th sample was retained. The polytope sampling algorithm used is the Gibbs sampler.

### 6.9 Sample diagnostics

An example of sampling diagnostics is shown in Figure 1, bottom left. This shows the autocorrelation functions for the two chains, for the flux from primary production to benthos (PP_Benthos) in the year 2011. This is obtained with the following command:

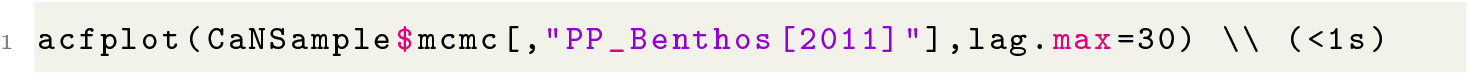

### 6.10 Sample Graphical outputs

An example of time-series outputs for 3 fluxes and 3 components is provided in Figure 1, bottom right. This is obtained with the following command:

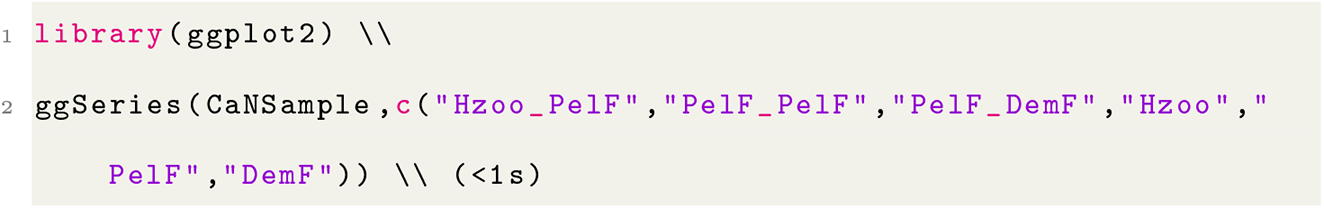

In addition to the R function ggSeries, RCaN provides a collection of predefined plotting functions which can be used to interpret the model outputs. In the Supplementary Information file, we provide examples of graphical outputs from these functions.

## Code and Data archiving statement

The code of RCaN and RCaNconstructor is available on github at https://github.com/inrae/RCaN. The code is published under a BSD3 license.

## Acknowledgements

The authors would like to thank Bérengère Husson, Elliot Sivel and Aurélien Favreau for their help in testing earlier version of the RCaN and RCaNconstructor softwares.

## Supplementary Information

A html file with a full description of the RCaN model implementation for the Barents Sea is provided as supplementary material. The Rmarkdown code and data sources used to generate the html file are available from the GitHub repository https://github.com/inrae/RCaN (folder article_supporting/PaperRCaN).

## A Appendix: Syntax of RCaN constraints

Constraints are at the heart of the Chance and Necessity modelling. They relate fluxes and biomass on the the trophic network. The basis is a mass balance equation: the variation of biomass of a trophic compartment is the result of assimilated input fluxes plus import fluxes minus output fluxes due to predation, losses due to somatic maintenance and other export fluxes. As somatic maintenance is proportional to biomass, this constraint is a linear one. The parameters: assimilation efficiency, trophic maintenance are given for all compartments in the RCaN-file. All linear constraints on fluxes and biomass can be expressed as constraints on trophic fluxes only. See [1] for precise formulation.

### A.1 Implicit and explicit constraints

There are two types of constraints.

1. Standard or implicit constraints. These are part of any RCaN model by default. They are defined by the value of the parameters in the Components sheet of the RCaN-file. These constraints express that (a) biomass must be positive and above the refuge biomass, (b) fluxes between two compartments are always positive, (c) the sum of fluxes into a compartment, from time t to t+1, cannot exceed *B*(*t*) * *σ*, where *B*(*t*) is the biomass at time t and *σ* is the satiation parameter, (d) the biomass of a compartment at time t+1 cannot be greater than *B*(*t*)*exp*(*ρ*) or lower than *B*(*t*)*exp*(−*ρ*), where *ρ* is the inertia coefficient.
2. Explicit constraints. These are specific to individual RCaN models. They express specific knowledge about how different biomass, fluxes and observations are related, i.e. constrained by each other. These explicit constraints are formulated in the constraints sheet of the RCaN-file, and must be written according to specific syntax rules. In this section we provide details and examples of this syntax.

### A.2 Symbolic expression of explicit constraints

Constraints are symbolic expressions, i.e. equalities or inequalities. They relates elements of the model (trophospecies or fluxes) on the left side of the expression to observations or other elements of the model on the right side of the expression. To interpret these expressions in a numerical framework, RCaN uses the R interface to the library Symengine, a fast symbolic manipulation library in C++, [14].

### A.3 Explicit constraints, principles and examples

The rules to write constraints explicitly are as follows.

1. Constraints are written in the form of inequalities or equalities.
2. The left side of the (in)equality must contains a reference to one or several model components or fluxes.
3. The right side of the (in)equality can contain fixed values, components, fluxes, observational time-series.

In the following constraints: *spA* and *spB* are the names (in the RCaN file) of two components, *fluxA_B* is the name of the flux between the two components, *obsA* is the name of an observational time-series of species *spA*. Some examples of standard constraints:

1. *spA* <= 100 the biomass of species A must be lower or equal to 100.
2. *spA* + *spB* <= 100 the combined biomasses of species A an B must be lower or equal to 100.
3. *fluxA_B* <= 50 the flux from species A to species B must be lower or equal to 50.
4. *spA* = *obsA* the biomass of species A must equate the observational time series of species A
5. *spA* <= *spB*: the biomass of species A must be lower or equal to the biomass of speciesB

### A.4 Explicit constraints using timeless absolute bounds

This type of constraint is useful when there are no data series available to inform on the temporal variations in certain biomass or fluxes but there is some knowledge about the maximum or minimum values that a compartment or a flux may take. For example, if the total biomass of species A (*spA*) is expected to lie between 100 and 1000 tonnes for the whole time series, one can write the following constraints:

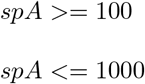

### A.5 Explicit constraints from imprecise biomass observations

Constraints using biomass time series, absolute estimates When times-series of absolute biomass estimates are available (e.g. from stock assessments) these can be used to constraint the corresponding modelled biomass. Consider a compartment inside the model (*spA*); a series of absolute biomass estimate with name *obsA*.

1. If we assume the observation to precisely reflect the true biomass, we can write the model constraint:

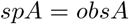
2. If we assume that the observed biomass is uncertain by 10% we can write the 2 model constraints:

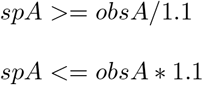
3. If we assume that the observed biomass only represent a fraction of the population and the the true biomass lies somewhere between what is estimated and twice this amount, we can write:

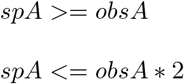

### A.6 Explicit constraints on Fisheries catches

The specific case of fisheries catches can be handled in the following way. Consider a species *A* (*spA*); a fishery on species *A* (compartment outside the model) with name *FA*. Catches are a flux from the species to the fishery (non-trophic flux): *spA_FA*. There is a time series of reported catches of species *A* by the fishery (data series); its name is *CatchA*. All are expressed in the same units (e.g. tonnes).

1. If we assume that the reported catches reflect the true catches exactly, we can write the model constraint:

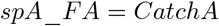
2. If we assume that the reported catches are uncertain by 10% we can write the model constraints:

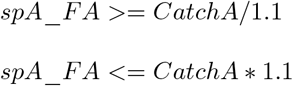
3. If we assume that the catches are under-reported and that the true catches are somewhere between what is reported and twice this amount, we can write:

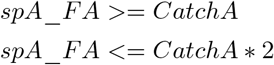

### A.7 Explicit constraints from relative estimates

When times-series of relative biomass estimates are available (e.g. from surveys) these can be used to constraint the corresponding modelled biomass. Consider compartment *A* (*spA*); a series of relative biomass estimate (data series) with name *surveyA*.

1. If we assume the observation to precisely reflect the relative variations in biomass over time, we can write the model constraint:

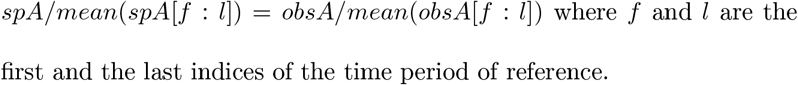
2. If we assume that the observed relative changes in biomass are uncertain by 10% we can write the 2 model constraints:

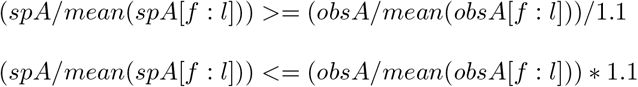

### A.8 Explicit structural constraints

In addition to constraining compartments/fluxes based on data or absolute bounds, it is also possible to express structural constraints within the model that are independent of observational time series or absolute bounds. For example, if species C (*spC*) can feed on species A and B (*spA* and *spB*), but we know that species A is always more abundant in the diet of species C, we can write.

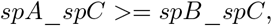

which expresses that the flux from *spA* to *spC* is always greater than the flux from *spB* to *spC*.

### A.9 Applying constraints over limited time periods

A time period is associated with each constraint. By default this is the period from the first year of available observation to the last. However, constraints can be applied to restricted time periods or specific years (or even a single year) when necessary. The selected years are indicated in the third column of the constraint table in the RCaN-constructor / RCaN-file.

