## Supplementary material for "RCaN : a software for Chance and Necessity modelling": barents_SM_RMD.html

Supplementary Information I : An application of CaN modelling to the Barents Sea trophic network


### Supplementary Information I : An application of CaN modelling to the Barents Sea trophic network

#### 5/2/2021

### 1 Introduction

We present here an example of RCaN study for the Barents Sea ecosystem. The CaN model has the same structure as the model presented in the main text of the article: same trophic structure, same trophic parameters, same set of time series of observations (Lindstrøm, Planque, and Subbey (2017)). Due to data access restriction, the values in the observation times series were slightly modified and do not correspond to the original measurements. Most of the original data can be provided on demand.

The network is illustrated in figure 1.1.

```
knitr::include_graphics("ReseauTrophiquePNG.png")
```

Figure 1.1: The Barents Sea tropic network. The dashed-line rectangle separate components that are within the model domain (herbivorous and omnivorous zooplankton, benthos, pelagic and demersal fish, mammals and birds) from those that are outside (primary producers, Norwegian Sea zooplankton and fisheries)

The RCaN-file, constructed with RCaNconstructor and the R-Markdown file, used to generate this supplementary material can be found on github.

We first provide information for the installation of RCaN and RCaNconstructor.
We then provide a short description of the content of the Barents Sea RCaN-file.
Finally, we go through the main steps of the modelling which follow the RCaN file construction: (a) build the polytope, (b) check the polytope status, (c) revise the model by adding and removing constraints, (d) sample the polytope, (d) analyze the samples.

### 2 Installing RCaN and RCaNconstructor

RCaN requires R (version 3.2+) and Rstudio to be installed.

The R library RCaN can be downloaded from https://github.com/inrae/RCaN. This is done by following installation instructions given in the ReadMe file of the package .

```
# if (!require("RCaN")) {
#  require(devtools)
#  devtools::install_git(
#    "https://gitlab.irstea.fr/hilaire.drouineau/can.git",
#    subdir="RCaN")
#  }
```

Then, the library can be loaded

```
library(RCaN) #the RCaN package itself
```

#### 2.1 Installing RCaNConstructor

- RCaNconstructor requires R (version 3.2+).
- RCaNconstructor requires the RCaN library to be installed;
- the RCaNconstructor installation procedure is described at https://github.com/inrae/RCaN/blob/master/RCaNconstructor/README.md

### 3 The RCaN file for the Barents Sea ecosystem

The food web structure, biological parameters, observational data-series and model constraints are all provided in the RCaN file in the .xlsx format. An RCaN file can be edited using a spreadsheet editor (e.g. libre office, open office) or with the RCaNconstructor. The use of RCaNconstructor is recommended as it ensures that names of model entities (species, fluxes and observations) are consistent across the different worksheets of the RCaN-file.

```
NAMEFILE <- 'BarentsSeaDummy.xlsx'
```

Importing an RCaN-file in the R environment and building the corresponding polytope are done with one single RCaN command:

```
CaNmod <- buildCaN(NAMEFILE)
```

The buildCaN function reads four worksheets from the RCaN-file and uses the information to construct the model, i.e. build the corresponding polytope. The information in these four worksheets pertains to components, fluxes, observational data-series and constraints. These are as follows:

##### 3.0.1 Components

Components correspond to the tropho-species in the systems (figure 1.1). They are specified in the “Components & input parameter” of the RCaN file, with their name (column Component), a flag to indicate whether they are inside or outside the model domain (column Inside) and their biological parameters. The biological parameters are used in the CaN master equation (which expresses mass conservation and the relationship between fluxes and biomass) and to implement implicit constraints (inertia, satiation and refuge biomass).

The above information is stored in the dataframe ‘components\_param’ in the CaNmod object:

```
datatable(CaNmod$components_param[c(1:8)], caption='Components')
```

In the Barents Sea example, the food-web has 10 components, 7 of which are within the model domain.

##### 3.0.2 Fluxes

Fluxes correspond to the transfer of biomass between food-web components (i.e., arrows in figure 1.1). They are specified in the “Fluxes” worksheet of the RCaN-file, identified by a name (column Flux), a source component (column From) and a sink component (column To) and a flag that indicates whether the flux is trophic or not (column Trophic). Non-trophic fluxes are typically import/export to/from the model domain and fisheries. The fluxes information is stored in the dataframe ‘flux\_def’ in the CaNmod object.

```
datatable(CaNmod$fluxes_def, caption='Fluxes')
```

In the Barents Sea example, the food-web includes 24 fluxes, 18 of which corresponding to trophic fluxes.

#### 3.1 Observational data-series

Observational time-series in the Barents Sea are derived from monitoring programs, mainly for plankton and fish species. In this example, the original data compiled by the ICES Working Group on the Integrated Ecosystem Assessment for the Barents Sea (ICES (2020)) have been altered to comply with data access restrictions (not all data is publicly available). They are stored in the “Input time-series” sheet in the RCaN-file. Each row indicate a time-step (year) and each column correspond to a particular data-series. The first row is a header with the name of each date-series. The observational data-series information is stored in the dataframe ‘series’ in the CaNmod object:

```
datatable(CaNmod$series,caption="Observations")
```

In the Barents Sea example, there are 22 time series of observations with a maximum length of 32 years.

#### 3.2 Constraints

Explicit constraints that relate food-web components, fluxes and observations can be specificied in the RCaN file. These are written in the form of inequalities or equalities and are provided in the worksheet “Constraints.” A constraint can be set to apply to every year or to selected years only. A constraint can be set to active/inactive (column ‘active’), which is practical for testing different combination of constraints without the need for writing/deleting the constraints in the worksheet. The constraint information is stored in the dataframe ‘constraints’ in the CaNmod object.

```
datatable(CaNmod$constraints[,1:5],caption='Constraints defined by user')
```

In the Barents Sea example, the model includes 36 constraints, 35 of them are active.

#### 3.3 Exploring the RCaN file with RCaNConstructor

The Barents Sea RCaN-file can be opened and explored directly in the RCaNconstructor. This is done with the following menus

- File - Open (alternatively “File - New” for creating a new model)
- Views - Network (graphically add, remove or edit components and fluxes)
- Views - Components (add, edit or remove components and their biological parameters)
- Views - Fluxes (add, edit or remove fluxes)
- Views - Observations (add, edit or remove observations, while specifying meta information)
- Views - Constraints (add, edit or remove constraints)

Figure 3.1: The Barents Sea food-web viewed from the graphical editor of RCaNconstructor. The dark-blue rectangle illustrates the model domain. Components within the model domain are shown in green and components outside the model domain are shown in red. Trophic links are shown in brown and non-trophic links are shown in green. Trophic links that go to and from the same component (cannibalism) are visualised by a thick circle around the correponding components.

### 4 The polytope behind a CaNmod object

In section 3, we used to the `buildCaN` function to read the RCaN file, but also to construct the corresponding polytope. The polytope is a multidimensional geometrical object that corresponds to the “space of possibles food-web trajectories.” The dimension of the polytope is defined by the number of fluxes times the number of time-steps (years) + the number of components. The different elements of the CaNmod object constructed using the buildCaN function are as follows:

```
names(CaNmod)
```

```
##  [1] "components_param" "species"          "fluxes_def"       "flow"            
##  [5] "series"           "ntstep"           "data_series_name" "constraints"     
##  [9] "H"                "N"                "A"                "AAll"            
## [13] "C"                "CAll"             "v"                "vAll"            
## [17] "L"                "b"                "bAll"             "symbolic_enviro"
```

Matrices C and A, and vectors b and v correspond to the matrices of active constraints that define the polytope A X <= b and C X = v (AAll, CAll, bAll and vAll are similar but include all constraints, active or not). Matrices N, H, F and L are matrices that are used to describe the dynamics of the system, more specifically, they describe how biomasses at any time can be derived from flow, using the relationship. The detailed mathematical formulation of the CaN model are provided in the supplementary material of Planque and Mullon Planque and Mullon (2019).

#### 4.1 Structure of the polytope

The polytope is defined inside a space with a high number of dimensions.

For the Barents sea example, we have:

```
paste("number of inequalities:", nrow(CaNmod$A))
```

```
## [1] "number of inequalities: 2593"
```

```
paste("number of equalities:", nrow(CaNmod$C))
```

```
## [1] "number of equalities: 64"
```

```
paste("number of parameters (dimensions):", ncol(CaNmod$C))
```

```
## [1] "number of parameters (dimensions): 775"
```

The inequalities and equalities result from both implicit and explicit constraints, each line corresponds to a constraint and a specific year. The number of parameters is the dimension of the polytope.

#### 4.2 Suitability of the polytope

The polytope contains the set of possible food-web trajectories. If some constraints are incompatible, the polytope will be empty, signifying that it is not possible to find a single food-web trajectory that can simultaneously satisfy all constraints. When there are too few constraints, the polytope can be unbounded. In such case, some of the fluxes can take an unbounded range of values, a somehow unrealistic situation.
Checking the polytope is an important step. Users can adopt different strategy: some users try first to specificy as many constraints as possible with the risk of constructing an empty polytope. They then need to relax some of the model constraints in order to find posible solutions. Others tend to start with a reduced set of constraints with the risk of having an unbounded polytope. They then need to add extra constraints to derive possible and plausible food-web trajectories. Both approaches have their pros and cons. Here, we focus on how to identify potential issues with the polytope and the possible ways to resolve them so that the polytope is suitable for sampling.

##### 4.2.1 First step: checking the polytope status

The R command *checkPolytopeStatus* returns the current status of the polytope. In the Barents Sea example, the polytope is ‘OK’ which means that it is non-empty and bounded. In other words, there is a limited set of possible trajectories.

```
checkPolytopeStatus(CaNmod)
```

```
## [1] "polytope ok"
```

In some situations, the function may return ‘empty polytope’ or ‘unbounded polytope.’ It might then be necessary to add/activate new constraints or remove/disactivate existing constraints.)

##### 4.2.2 Toggling constraints

Constraints can be activated/deactivate in the RCaN-file, using the ‘active’ switch in the ‘Constraints’ worksheet. In addition, it is possible to activate/deactivate constraints directly within R, once the RCaN file has been loaded. We illustrate this below, using the command *toggleConstraint*:

```
CaNmod <- toggleConstraint(CaNmod,
                        c("C01","C02","C03","C04","C05","C06","C07","C08","C09","C10","C11","C12","C15","C16","C19","C21"))
```

```
...
##  [1] "disactivate inequality C01 : 1998" "disactivate inequality C01 : 1999"
##  [3] "disactivate inequality C01 : 2000" "disactivate inequality C01 : 2001"
##  [5] "disactivate inequality C01 : 2002" "disactivate inequality C01 : 2003"
##  [7] "disactivate inequality C01 : 2004" "disactivate inequality C01 : 2005"
##  [9] "disactivate inequality C01 : 2006" "disactivate inequality C01 : 2007"
## [11] "disactivate inequality C01 : 2008" "disactivate inequality C01 : 2009"
## [13] "disactivate inequality C01 : 2010" "disactivate inequality C01 : 2011"
## [15] "disactivate inequality C01 : 2012" "disactivate inequality C01 : 2013"
## [17] "disactivate inequality C01 : 2014" "disactivate inequality C01 : 2015"
## [19] "disactivate inequality C01 : 2016" "disactivate inequality C01 : 2017"
...
```

In this example, we removed constraints C01 to C12 and C15, C16, C19 and C21, which bound the input (primary production), the output of the system (fisheries) and the maximum biomass of top predators (birds and marine mammals).

```
checkPolytopeStatus(CaNmod)
```

```
## [1] "polytope not bounded"
```

As a result, the polytope is now unbounded.
We therefore reactivate the constraint with the ‘toggleConstraint’ command.

```
CaNmod <- toggleConstraint(CaNmod,
                        c("C01","C02","C03","C04","C05","C06","C07","C08","C09","C10","C11","C12","C15","C16","C19","C21"))
```

```
...
##  [1] "activate inequality C01 : 1998" "activate inequality C01 : 1999"
##  [3] "activate inequality C01 : 2000" "activate inequality C01 : 2001"
##  [5] "activate inequality C01 : 2002" "activate inequality C01 : 2003"
##  [7] "activate inequality C01 : 2004" "activate inequality C01 : 2005"
##  [9] "activate inequality C01 : 2006" "activate inequality C01 : 2007"
## [11] "activate inequality C01 : 2008" "activate inequality C01 : 2009"
## [13] "activate inequality C01 : 2010" "activate inequality C01 : 2011"
## [15] "activate inequality C01 : 2012" "activate inequality C01 : 2013"
## [17] "activate inequality C01 : 2014" "activate inequality C01 : 2015"
## [19] "activate inequality C01 : 2016" "activate inequality C01 : 2017"
...
```

##### 4.2.3 Finding incompatible constraints

We artificially add new constraint which is incompatible with the existing ones to illustrate a situation where the polytope is empty.

```
CaNmod<-toggleConstraint(CaNmod, "C35")
```

```
...
##  [1] "activate inequality C35 : 1988" "activate inequality C35 : 1989"
##  [3] "activate inequality C35 : 1990" "activate inequality C35 : 1991"
##  [5] "activate inequality C35 : 1992" "activate inequality C35 : 1993"
##  [7] "activate inequality C35 : 1994" "activate inequality C35 : 1995"
##  [9] "activate inequality C35 : 1996" "activate inequality C35 : 1997"
## [11] "activate inequality C35 : 1998" "activate inequality C35 : 1999"
## [13] "activate inequality C35 : 2000" "activate inequality C35 : 2001"
## [15] "activate inequality C35 : 2002" "activate inequality C35 : 2003"
## [17] "activate inequality C35 : 2004" "activate inequality C35 : 2005"
## [19] "activate inequality C35 : 2006" "activate inequality C35 : 2007"
...
```

From section 3.2, we can see that constraint C35 is not incompatible with constraints C21, i.e. the two can not be fulfilled simultaneously. As a consequence, checking the polytope status returns ‘empty polytope’:

```
checkPolytopeStatus(CaNmod)
```

```
## [1] "empty polytope"
```

The RCaN function ‘findingIncompatibleConstr’ is then useful to identify which constraints are incompatible:

```
incomp <- findingIncompatibleConstr(CaNmod)
```

```
...
## [1] "###polytope is ok when following constraints are relaxed:"
## [1] " C35 : 1988,  C35 : 1989,  C35 : 1990,  C35 : 1991,  C35 : 1992,  C35 : 1993,  C35 : 1994,  C35 : 1995,  C35 : 1996,  C35 : 1997,  C35 : 1998,  C35 : 1999,  C35 : 2000,  C35 : 2001,  C35 : 2002,  C35 : 2003,  C35 : 2004,  C35 : 2005,  C35 : 2006,  C35 : 2007,  C35 : 2008,  C35 : 2009,  C35 : 2010,  C35 : 2011,  C35 : 2012,  C35 : 2013,  C35 : 2014,  C35 : 2015,  C35 : 2016,  C35 : 2017,  C35 : 2018,  C35 : 2019"
## [1] "####Those constraints seem incompatible with:"
## [[1]]
## [1] " C35 : 1988:  C27b : 2001,  C29 : 2016,  C29 : 2017,  C29 : 2018,  C29 : 2019,  C34 : 1988,  C34 : 1989,  C34 : 1990,  C34 : 1991,  C34 : 1992,  C34 : 1993,  C34 : 1994,  C34 : 1995,  C34 : 1996,  C34 : 1997,  C34 : 1998,  C34 : 1999,  C34 : 2000,  C34 : 2001,  C34 : 2002,  C34 : 2003,  C34 : 2004,  C34 : 2005,  C34 : 2006,  C34 : 2007,  C34 : 2008,  C34 : 2009,  C34 : 2010,  C34 : 2011,  C34 : 2012,  C34 : 2013,  C34 : 2014"
## 
## [[2]]
## [1] " C35 : 1989:  C27b : 2016,  C29 : 2018,  C29 : 2019,  C34 : 1988,  C34 : 1989,  C34 : 1990,  C34 : 1991,  C34 : 1992,  C34 : 1993,  C34 : 1994,  C34 : 1995,  C34 : 1996,  C34 : 1997,  C34 : 1998,  C34 : 1999,  C34 : 2000,  C34 : 2001,  C34 : 2002,  C34 : 2003,  C34 : 2004,  C34 : 2005,  C34 : 2006,  C34 : 2007,  C34 : 2008,  C34 : 2009,  C34 : 2010,  C34 : 2011,  C34 : 2012,  C34 : 2013,  C34 : 2014,  C34 : 2015,  C34 : 2016"
## 
## [[3]]
...
```

The function returns that constraint C35 is problematic (see ###polytope is ok when following constraints are relaxed) and that it is incompatible with constraints C16 and C17 (“####Those constraints seem incompatible with:”).

In such situation, it is necessary to relax some of the above constraints (C35, C16, C17) to obtain a non-empty polytope. The choice of the constraint to relax is left to the modeller, based on domain knowledge.

In the following, we revert to a model version in which constraint C35 is inactive and the polytope is OK:

```
CaNmod<-toggleConstraint(CaNmod, "C35")
```

```
...
##  [1] "disactivate inequality C35 : 1988" "disactivate inequality C35 : 1989"
##  [3] "disactivate inequality C35 : 1990" "disactivate inequality C35 : 1991"
##  [5] "disactivate inequality C35 : 1992" "disactivate inequality C35 : 1993"
##  [7] "disactivate inequality C35 : 1994" "disactivate inequality C35 : 1995"
##  [9] "disactivate inequality C35 : 1996" "disactivate inequality C35 : 1997"
## [11] "disactivate inequality C35 : 1998" "disactivate inequality C35 : 1999"
## [13] "disactivate inequality C35 : 2000" "disactivate inequality C35 : 2001"
## [15] "disactivate inequality C35 : 2002" "disactivate inequality C35 : 2003"
## [17] "disactivate inequality C35 : 2004" "disactivate inequality C35 : 2005"
## [19] "disactivate inequality C35 : 2006" "disactivate inequality C35 : 2007"
...
```

#### 4.3 Bounds of the polytope

To obtain a first overview of the polytope, it is useful to compute its limits in all dimensions. This can be achieved with the RCaN function *getAllBoundsParam*. The output is a dataframe with a row per parameter/dimension. It is time consuming to compute all bounds, and one alternative is to compute the bounds for a small set of dimensions only. This can be done using the function *getBoundParam*. This is illustrated below for the two dimensions corresponding to the flux from Omnivorous Zooplankton to Birds in 2017 and 2018:

```
getBoundParam(CaNmod, which(colnames(CaNmod$A)=="Ozoo_Birds[2017]"))
```

```
## [1]    0.000 2666.667
```

```
getBoundParam(CaNmod, which(colnames(CaNmod$A)=="Ozoo_Birds[2018]"))
```

```
## [1]    0.000 2666.667
```

#### 4.4 2D-slices of the polytope

2-dimensional pictures of the polytope are useful to visualise how the bounds of different fluxes are jointly constrained. These are obtained using the RCaN function *plotPolytope2D*:

```
start_time <- Sys.time()
fluxX1 <- 'PP_Ozoo[1990]'
fluxY1 <- 'Hzoo_Ozoo[1990]'
plotPolytope2D(CaNmod, c(fluxX1, fluxY1), progressBar=FALSE)
```

```
fluxX2 <- 'Hzoo_PelF[1990]'
fluxY2 <- 'PelF_PelF[1990]'
plotPolytope2D(CaNmod, c(fluxX2, fluxY2), progressBar=FALSE)
```

```
fluxX3 <- 'Hzoo_PelF[1990]'
fluxY3 <- 'Hzoo_PelF[1991]'
plotPolytope2D(CaNmod, c(fluxX3, fluxY3), progressBar=FALSE)
```

```
fluxX4 <- 'Hzoo_PelF[1990]'
fluxY4 <- 'PelF_Fisheries[1991]'
plotPolytope2D(CaNmod, c(fluxX4, fluxY4), progressBar=FALSE)
```

```
end_time <- Sys.time()
end_time - start_time
```

```
## Time difference of 10.54574 mins
```

#### 4.5 Doing the same with RCaNConstructor

- RCaN - Polytope - Build polytope
- RCaN - Views - Constraints : edit the Active parameter, add or remvove constraints
- RCaN - Polytope - Analyze polytope - Plot network
- RCaN - Polytope - Analyze polytope - Diagnostic polytope
- RCaN - Polytope - Analyze polytope - Find incompatible constraints
- RCaN - Polytope - Analyze polytope - Plot polytope in two selected dimensions

### 5 Sampling the polytope

#### 5.1 Sampling

The RCaN function *sampleCaN* is used to sample of the polytope, i.e. to sample several random possible trajectories of the food-web dynamics. The output of this command is a mcmc object (defined in package coda). The function parameters include:
\* N: the number of samples
\* thin: the thinning, i.e. the number of samples that are left out from the output
\* nchains: the number of sampling chains
\* ncore: the number of CPU cores used
\* method: one of gibbs (default) or hitandrun

With N=100, thin=10, and nchain=2 the *sampleCaN* function will sample two chains of 1,000 values and return one output every 10, that is a total of 200 valid samples

```
system.time(samples <- sampleCaN(CaNmod, N=100, thin=10, nchain=2, ncore=2))
```

```
## utilisateur     système      écoulé 
##       0.123       0.017     199.102
```

#### 5.2 Convergence of the sample chains

Check the mixing of the Monte Carlo Marko chain algorithm that is used to build the sample can be achieved with commands from the coda library.

```
summ <- summary(samples$mcmc)

datatable( cbind.data.frame(summ$statistics, summ$quantiles) %>%
             select('Mean', 'SD', '2.5%', '50%', '97.5%'),
           options = list(pageLength = 31)) %>%
  formatRound(attr(.$x,"colnames"))
```

##### 5.2.1 Gelman diagnostic

Gelman and Rubin (1992) proposed a general approach to asses the convergence of MCMC outputs in which $ m > 1$ parallel chains are run with starting values that are overdispersed relative to the posterior distribution. Convergence is diagnosed when the chains have ‘forgotten’ their initial values, and the output from all chains is indistinguishable. The gelman.diag diagnostic is applied to a single variable from the chain. It is based a comparison of within-chain and between-chain variances, and is similar to a classical analysis of variance.

```
fluxZ1 <- 'PP_Benthos[1990]'
fluxZ2 <- 'Ozoo_Birds[1990]'
fluxZ3 <- 'DemF_Fisheries[1990]'
fluxZ4 <- 'PelF_DemF[1990]'
gelman.diag(samples$mcmc[,c(fluxZ1,fluxZ2,fluxZ3,fluxZ4)], multivariate = FALSE)
```

```
## Potential scale reduction factors:
## 
##                      Point est. Upper C.I.
## PP_Benthos[1990]           1.03       1.06
## Ozoo_Birds[1990]           1.05       1.24
## DemF_Fisheries[1990]       1.04       1.21
## PelF_DemF[1990]            1.04       1.18
```

Theoretically a statistics below 1.05 suggests an appropriate convergence.

##### 5.2.2 Autocorrelation function

The *acfplot* function from the coda library shows the autocorrelation function of sampling chains. It is illustrative of the independence of successive samples. If the samples are not serially correlated (which is desired) the autocorrelation should remain around zero (except for the first value which is always 1). Otherwise, it might be worthwhile to increase the thinning during RCaN sampling and/or to discard the first samples.

```
fluxZ1 <- 'PP_Benthos[1990]'
fluxZ2 <- 'Ozoo_Birds[1990]'
acfplot(samples$mcmc[,fluxZ1], ylim=c(-0.1,0.8), main=fluxZ1)
```

```
acfplot(samples$mcmc[,fluxZ2], ylim=c(-0.1,0.8), main=fluxZ2)
```

### 6 Visualisation of the model results

#### 6.1 Temporal dynamics of componentns and fluxes

The graphical representation of the possible dynamics of the food-web is a important output of the model. The RCaN function *ggSeries* produces standardised plots of the time-series of components and/or fluxes. By default, for each component, 3 randomly picked samples/trajectories are plotted (plain, dashed and dotted lines) together with the envelopes containing 100%, 95% and 50% of the sampled trajectories.

For components, we get:

```
ggSeries(samples, CaNmod$species, TRUE) +
  theme(legend.position = "none")
```

In this example we observe different situations for the different species: for pelagic fish (PelF) and demersal fish (DemF) the constraints imposed in relation to the observational data-series constrain the trajectories of biomass which are all situated within a narrow band. On the other hand, there is little information provided to constriain directly the trajectories of marine mammal and Birds which leads to large variability between the different samples.

The dynamics of fluxes can be plotted in a similar fashion:

```
ggSeries(samples, CaNmod$flow, TRUE) +
  theme(legend.position = "none")
```

#### 6.2 Empirical distributions of biomass and fluxes

Exploring the distribution of biomass and fluxes across samples and across years is also a useful output of the model. The RCaN function *ggViolin* produces standardised violin plots of these distributions. By default, these are presented on a log scale which allows for comparison between biomass or fluxes that can span several orders of magnitude. For example, the distribution of species biomass for the year 1990 is obtained as follows:

```
ggViolin(samples,CaNmod$species,year=1990,TRUE) + 
  xlab('Component') + ylab('Biomass')
```

The violin plots of all fluxes in 1990 are produced as follows::

```
ggViolin(samples,CaNmod$flow,year=1990,TRUE) + 
  xlab('Component') + ylab('Biomass') +
  theme(axis.text.x=element_text(angle=90))
```

#### 6.3 Diet composition

The diet matrix is a key parameter of many trophic models, such as e.g. Ecopath (Christensen and Walters (2004)). In CaN models, diet is non deterministic, in the sense that trophic fluxes are constrained but not determined by biomass. Diets are thus emerging from the model and it is interesting to explore the diets of individual species. A convenient way is to use the RCaN function *ggDiet* to draw the average diet for each predators:

```
ggDiet(samples, CaNmod$species) + 
  ylab('prey proportion in the diet')
```

#### 6.4 Growth and density-dependence

Density-dependence usually reflects how growth is related to population size In RCaN trajectories this can be expressed as growth in biomass in relation to biomass. The RCaN function *ggGrowth* plots the growth rate (measured as the ratio of biomass at time $ t+1 $ over the biomass at time $ t $) as a function of the biomass (at time $ t $). Plotting density-dependence for all species is done as follows:

```
ggGrowth(samples, CaNmod$species)
```

This illustrates how density-dependence of the food-web components can emerge from the food-web dynamics. The upper and lower red-dotted lines visualise the minimum and maximum growth rates set by the inertia constraints for each component. The thick lines are smoothers which highlight the possible underlying relationship between growth and biomass.

#### 6.5 Trophic functional relationships

The RCaN function *ggTrophicRelation* plots the empirical trophic functional relationships (TFRs) between the biomass of a prey and the quantity of that prey consumed by a predator. Below is the command for plotting the TFR between benthos, pelagic fishes and demersal fishes in the Barents Sea example.

```
ggTrophicRelation(samples, CaNmod$species[3:5])
```

Above, we see that the predation of benthos by demersal fishes does not increases with the biomass of benthos, suggesting that benthos is not a limiting factor in the diet of demersal fishes.

#### 6.6 Satiation

The function *ggSatiation* plots the total biomass of prey eaten by a predator, as a function of the biomass of the predator. In addition the plot includes an indication (upper red dotted line) of the maximum possible consumption rate, derived from the satiation parameter of the predator species. An example is illustrated below for all model components and over the entire sampling period. It shows that marine mammals never reach their satiation. On the other hand, demersal fish are often closed to satiation, suggesting that they are not food limited.

```
ggSatiation(samples, CaNmod$species)
```

#### 6.7 Feeding and Growth

The RCaN function *ggSatiatInertia* is used to explore the relationships between population growth and feeding. The plot shows standardized total consumption(0=no feeding, 1=feeding to satiation) as a function of standardized population growth/mortality (-1 = maximum mortality, 0 = no change in biomass, 1 = maximum growth). A positive relationship between growth and satiation is indicative of bottom up control.

```
ggSatiatInertia(samples, CaNmod$species)
```

#### 6.8 Pair-plots

The function *ggPairsBiomass* is used to explore the relationships between the biomass of different food-web components/species. The plot shows the individual component density function in the diagonal, the scatterplots of one species against the other in the lower triangle and the Kendall correlation coefficient in the upper triangle. An example is illustrated below for three model components and over the entire sampling period. The function parameter *frac* indicates the fraction of the samples which will be plotted. This is useful for large sample size which can lead to overloaded figures.

```
ggPairsBiomass(samples,c("Benthos","PelF","DemF"),logscale=FALSE, frac = .25) + #we only 25% of the point since the function is long
  scale_y_continuous(n.breaks=3)+ scale_x_continuous(breaks=c(1990,2010))
```

#### 6.9 Top-Down and Bottom-Up controls

One important aspect of food-web dynamics is the presence of possible trophic controls (Bottom-up or Top-Down). We define here bottom-up as a situation in which the growth of a species/component is positively correlated to the consumption by this species. Similarly, Top-down control is defined as a situation in which the growth of a species/component is negatively correlated to the predation/fishing on this species. The RCaN function *ggTopDownBottomUp* plots the density function of the correlation coefficients between growth and consumption/predation:

```
ggTopDownBottomUp(samples)
```

This function display the correlation between the growth of the species with both (1) its total ingestion of prey per unit of biomass (in red) and (2) the how much it is predated per unit of biomass (in green). A red density plot tending to 1 indicates that growth is correlated to feeding, and therefore a bottom up control. Conversely, a green density plot tending towards minus one indicates that growth is negatively correlated to the predation and suggests a top down control. It is possible to select which fluxes are accounted for in this diagram (for example to exclude non-trophic fluxes):

```
# we restrict the analysis to Ozoo and DemF and limit to the predation by
#respectively DemF/PelF and MM
g <- ggTopDownBottomUp(samples, species=list(Ozoo=c("DemF","PelF"), DemF=c("MM"))) 
g
```

#### 6.10 Doing the same with RCaNConstructor

- RCaN - Sample - Sample polytope
- RCaN - Sample - Analyze polytope - Gelman diagnostic
- RCaN - Sample - Analyze polytope - Plot autocorrelation of sample
- RCaN - Sample - Analyze polytope - Plot sample
- RCaN - Sample - Analyze polytope - Violin plots of biomass distributions
- RCaN - Sample - Analyze polytope - Violin plots of flux distributions
- RCaN - Sample - Analyze polytope - Diet fractions
- RCaN - Sample - Analyze polytope - Growth / Biomasses
- RCaN - Sample - Analyze polytope - Trophic functional relationships
- RCaN - Sample - Analyze polytope - Satiation
- RCaN - Sample - Analyze polytope - Satiation/Inertia
- RCaN - Sample - Analyze polytope - Pairs of biomass

Planque, Benjamin, and Christian Mullon. 2019. “Modelling Chance and Necessity in Natural Systems.” *ICES Journal of Marine Science*.
